# Sequence variation aware genome references and read mapping with the variation graph toolkit

**DOI:** 10.1101/234856

**Authors:** Erik Garrison, Jouni Sirén, Adam M. Novak, Glenn Hickey, Jordan M. Eizenga, Eric T. Dawson, William Jones, Michael F. Lin, Benedict Paten, Richard Durbin

## Abstract

Reference genomes guide our interpretation of DNA sequence data. However, conventional linear references are fundamentally limited in that they represent only one version of each locus, whereas the population may contain multiple variants. When the reference represents an individual’s genome poorly, it can impact read mapping and introduce bias. Variation graphs are bidirected DNA sequence graphs that compactly represent genetic variation, including large scale structural variation such as inversions and duplications.^1^ Equivalent structures are produced by de novo genome assemblers.^2,3^ Here we present vg, a toolkit of computational methods for creating, manipulating, and utilizing these structures as references at the scale of the human genome. vg provides an efficient approach to mapping reads onto arbitrary variation graphs using generalized compressed suffix arrays,^4^ with improved accuracy over alignment to a linear reference, creating data structures to support downstream variant calling and genotyping. These capabilities make using variation graphs as reference structures for DNA sequencing practical at the scale of vertebrate genomes, or at the topological complexity of new species assemblies.

## 1 Introduction

Where genomes are small, it is possible to study genetic variation by assembling whole genomes and then comparing them via whole-genome comparison.^5,6^ For large genomes such as human, routine complete and accurate *de novo* genome assembly is impractical, and we use prior information to interpret new sequence data in its correct genomic context. The current paradigm is to align sequence reads to a single high-quality reference genome sequence that represents one haplotype at each location in the genome, leading to mapping biases towards the reference and away from variant alleles. There will even be some sequence in each new sample that is entirely absent in the reference.^7^

In principle, we would like to align to a genome that is as similar to our sample as possible, ideally a “personalized” reference genome^8^ that already incorporates the individual’s variants. Although in general we will not know what variants are present before aligning data from a sample, most differences between any one genome and the reference are segregating in the population. Thus, if we build a structure that represents known shared variation, then that will contain most of the correct personalized reference sequence for any individual.

The natural computational structure for doing this is the sequence graph.^1^ Sequence graphs have long been used to represent multiple sequences that contain shared differences or ambiguities in a single structure. For example, multiple sequence alignments have a natural representation as partially ordered sequence graphs.^9^ The variant call format^10^ (VCF), which is a common data format for describing populations of genome sequences can be understood as defining a partially ordered graph similar to those implied by a multiple sequence alignment. Related structures frequently used in genome assembly include the De Bruijn graph^2^ and string graph,^3^ which collapse long repeated sequences, so the same nodes are used for different regions of the genome. Graphs to represent genetic variation have previously been used for microbial genomes and localized regions of the human genome such as the Major Histocompatibility Complex.^11^

We define a variation graph as a sequence graph together with a set of paths representing possible sequences from a population (Figure 1). In this paper, we describe vg, a multipurpose toolkit for performing genomic analyses using a variation graph as a reference, discussing in particular construction, modification and read mapping. Recently software packages have been introduced that support a subset of variation graphs that reflect local variation away from a linear reference,^12,13^ formalizing approaches introduced in FreeBayes and the GATK HaplotypeCaller for the 1000 Genomes Project analysis.^14-16^ Importantly, our model goes beyond these in that it does not require the graph to be based around an initial linear reference, or indeed directionally ordered, and thus supports cycles and inversions. vg is the first openly available tool with these properties to scale practically to the multi-gigabase scale required for whole vertebrate genomes.

**Figure 1:**
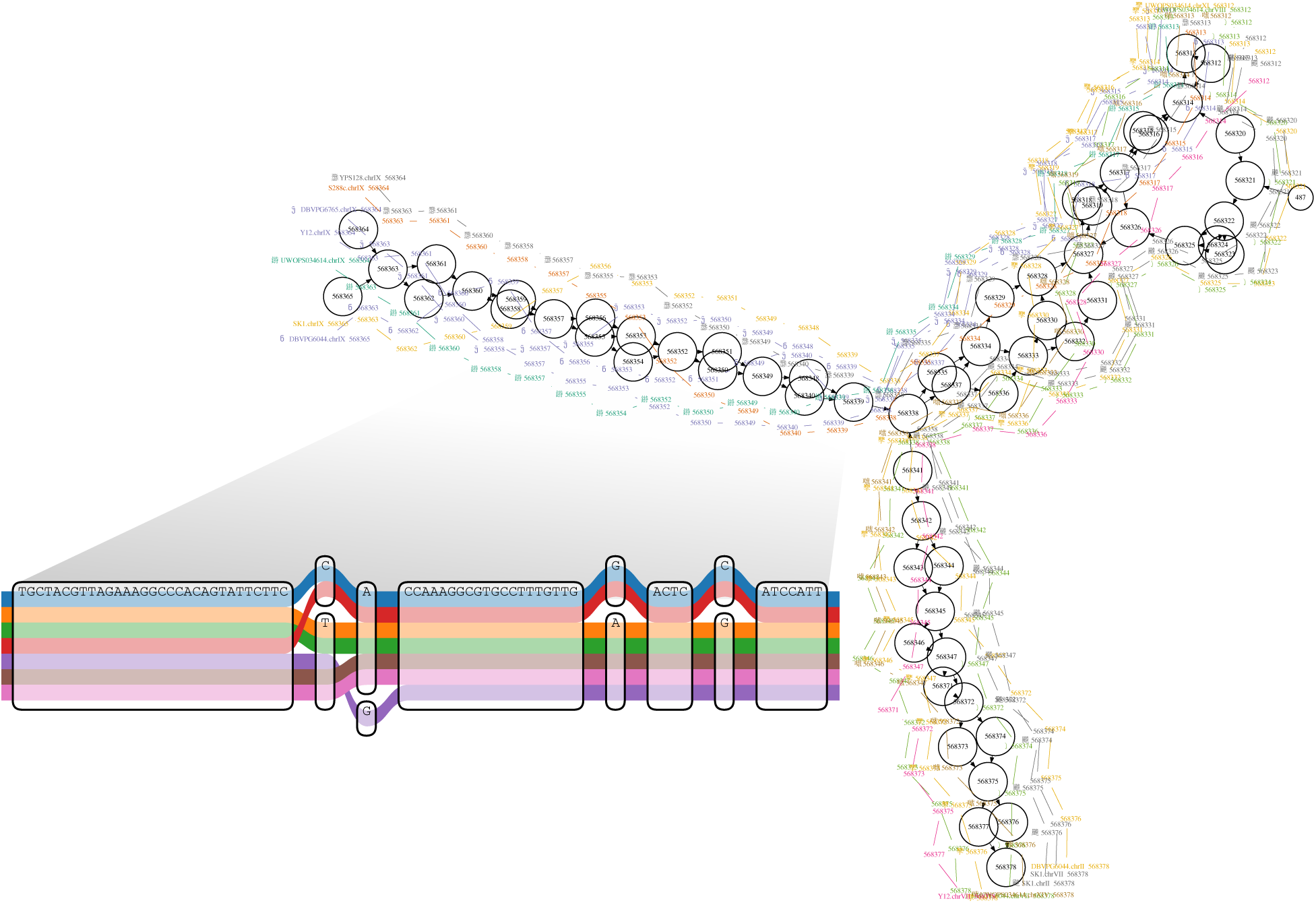
A region of a yeast genome variation graph corresponding to the start of the subtelomeric region on the left arm of chromosome 9 in a multiple alignment of the strains sequenced by Yue et al.,^18^ built using vg from a full genome multiple alignment generated with the Cactus alignment package.^6^ The inset shows a subregion of the alignment at single base level. The colored paths correspond to separate contiguous chromosomal segments of these strains.

Here we describe the design and implementation of vg, and demonstrate its capabilities by constructing a variety of variation graphs and aligning sequencing data against these, demonstrating that we observe improved mapping compared to linear references. We first describe the core data model, data structures and algorithms, and implementation of vg. We then we present results demonstrating the functionality of vg. Variant calling using vg against a variety of different human genome variation graphs is described elsewhere.^17^

## 2 Methods

### 2.1 Model

We define a variation graph to be a graph with embedded paths *G* = (*N,E,P*) comprised of a set of nodes *N* = *n*_1_… *n_M_*, a set of edges *E* = *e*_1_… *e_L_*, and a set of paths *P* = *p*_1_ … *p_Q_*, each of which describes the embedding of a sequence into the graph.

Each node *n_i_* represents a sequence *seq*(*n_i_*) which is built from an alphabet *A* = {A, C, G, T, N}. Nodes may be traversed in either the forward or reverse direction, with the sequence being reverse-complemented in the reverse direction. We write 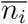 for the reverse-complement of node *n_i_*, so that 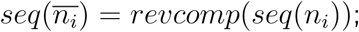 note that 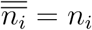. For convenience, we refer to both *n_i_* and 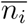 as “nodes”.

Edges represent adjacencies between the sequences of the nodes they connect. Thus, the graph implicitly encodes longer sequences as the concatenation of node sequences along walks through the graph. Edges can be identified with the ordered pairs of oriented nodes that they link, so we can write *e*_*i*→*j*_ = (*n_i_*,*n_j_*). Edges also can be traversed in either the forward or the reverse direction, with the reverse traversal defined as 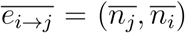. Note that graphs in vg can contain ordinary cycles (in which *n_i_* is reachable from *n_i_*), reversing cycles (in which 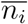 is reachable from *n_i_*), and non-cyclic instances of reversal (in which both *n_i_* and 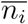 are reachable from *n_j_*).

### 2.2 Implementation

The vg implementation is multithreaded and written in C++11, and is available from http://github.com/vgteam/vg under the MIT open source software license. It provides both a primary application to support the operations we describe here, and a library libvg which applications can use to access the data structures, indexes and low level operations.

Our core representation of the graph uses Google’s open source protobuf system, which directly supports serialization onto disk for storage. We also provide a protobuf alignment format, GAM (for “Graph Alignment Map”), with analogous functionality to BAM.^19^ To enable read mapping and other random access operations against large sequence graphs we have implemented a succinct representation of a vg variation graph (xg) that is static but very memory and time efficient, using rank/select dictionaries and other data structures from the Succinct Data Structures Library (SDSL).^20^ Graphs can be imported from and exported to a variety of formats, including the assembly format GFA and the W3C graph exchange format RDF.^21^ Further details about the implementation and features are available in the supplement and at the Github website.

### 2.3 Alignment

A key requirement for a reference genome is the ability to efficiently and accurately find an optimal alignment for a new DNA sequence such as a sequencing read. Analogous to the way that read mappers to linear references work, our approach to this problem is to find seed matches by an indexed search process, cluster them if there are multiple seeds close together, and then perform a local constrained dynamic programming alignment of the read against a region of the graph around each cluster. A brief description of the key steps in this process is given here, with further details in the supplement.

The GCSA2 library^4^ that vg uses for seeding can perform linear time exact match queries independent of the graph size to find super-maximal exact match (SMEM) seeds, subject to a maximum query length, in time comparable to the corresponding operations in bwa mem. SMEMs are exact matches between a query substring and a reference substring that cannot be extended in either direction, and for which there is no extension of the query substring that matches elsewhere in the graph.

After obtaining SMEMs for a query sequence using GCSA2, we cluster them using a global approximate distance metric and distance estimates provided by any nearby paths. For paired reads, we cluster all the SMEMs for both reads in the pair to preferentially support mappings where the SMEMs match a fragment model that we establish online during the alignment of the read set.

We next select the best consistent sequence of SMEMs within each cluster as the maximum likelihood path through a Markov model over the SMEMs that rewards long SMEMs and short gaps between SMEMs. In many cases there is just one SMEM in the sequence, but there are complex cases where the best SMEM sequence is not correct, and to catch these we recursively mask out the SMEMs in paths found so far and re-run the algorithm to obtain additional disjoint SMEM sequences if available.

For each consistent sequence of SMEMs, we then obtain the subgraph containing the cluster. To avoid the complications introduced by cycles and inversions,^22^ we transform the local graph region into a directed acyclic graph (DAG) while maintaining an embedding in the original, potentially cyclic bidirected graph. We can then perform partial order alignment to the DAG,^9^ using banded dynamic programming and an extension of Farrar’s SIMD-accelerated striped Smith Waterman algorithm.^23^

The vg alignment tool also uses base qualities in alignment scores, and calculates adjusted mapping quality scores. Base qualities are probabilistic estimates of the confidence of each base call in a read provided by the sequencing technology. vg combines these with a probabilistic interpretation of alignment^24^ to adjust the scoring function for alignments, which has previously been shown to improve variant calling accuracy.^25^ Mapping qualities^26^ are a probability-based measure of the confidence in the localization of a read on the reference that is important for variant calling and other downstream analyses. vg computes mapping qualities by comparing the scores of optimal and subobptimal alignments under the probabilistic alignment model, in a similar fashion to bwa mem.

### 2.4 Graph editing and construction

We can build a graph either by direct construction from external graphs such as from de novo assemblies, or by a series of editing operations applied to simple starting graphs such as standard linear reference genomes. To support editing of existing graphs, vg supports operations that can split a node where sequences diverge and insert additional edges and nodes. While doing this it keeps track of the relationship to the previous graph in a *translation* object, which supports projection of coordinates from one version of the graph to another.

We make use of the editing operations to construct graphs from Variant Call Format (VCF) files,^10^ as produced by population sequencing projects such as the 1000 Genomes Project,^16^ inserting a cluster of nodes and edges into a linear reference for each overlapping subset of VCF records. Edit operations also allow progressive construction of a vg graph from a set of sequences by repeated alignment and editing, so that all the initial sequences are embedded in the graph as paths. Last but not least, edit operations allow new variants to be added to an existing vg reference graph to support use cases such as incorporating novel variants from new individuals mapped and called against the graph, while retaining a coordinate mapping to the existing reference. These actions are also invertible, in that vg can generate VCF to describe the graph as a set of variants, using an arbitrarily chosen embedded path as a reference.

### 2.5 Experiments

Experiments were carried out on a dedicated compute node with 256 gigabytes of RAM and two 2.4GHz AMD Opteron 6378 processors with a total of 32 CPU cores, or for the whole human data set on a cluster of Amazon Web Services r3.8xlarge nodes.

## 3 Results

For a species such as human with only 0.1% nucleotide divergence on average between individual genome sequences, over 90% of 100bp reads will derive from sequence exactly matching the reference. Therefore the first requirement for any new mapper to be used in practice is that against a linear reference it performs at least as well as the current standard, which we take to be bwa mem^27^ with default parameters. We then show that around divergent sites vg maps significantly more informatively.

### 3.1 Mapping human data to the 1000 Genomes Project graph

The final phase of the 1000 Genomes Project produced a data set of approximately 80 million variants in 2504 humans.^16^ The corresponding vg graph uses 4.76 GB when serialized to disk, and contains 3.181 Gbp of sequence, which is exactly equivalent to the length of the input reference plus the length of the novel alleles in the VCF file. The associated random-access and search indices require 38 GB memory each. We also built and indexed a second vg graph corresponding to the standard GRCh37 linear reference sequence, without including any variation.

We next aligned ten million 150 bp paired end reads simulated with errors from the parentally phased haplotypes of an Ashkenazi Jewish male NA24385, who was sequenced by the Genome in a Bottle (GIAB) Consortium,^28^ and who is not included in the 1000 Genomes Project sample set. To map the reads to the 1000GP graph required 60 seconds to load the indexes, 75 GB of RAM, and approximately one elapsed hour (32 core hours) to align the reads. Figures 2a and 2b show the accuracy of these alignments compared with bwa mem, in terms of Receiver Operating Characteristic (ROC) curves parameterised by mapping quality, partitioned into reads that were simulated from segments in N24385 not containing variants from the reference (Figure 2a), and reads simulated from segments that do contain variants (Figure 2b). Both groups of reads may contain differences from the source segment sequences because of simulated sequencing errors, which included both substitution and (at a lower rate) insertion/deletion (indel) errors.

**Figure 2:**
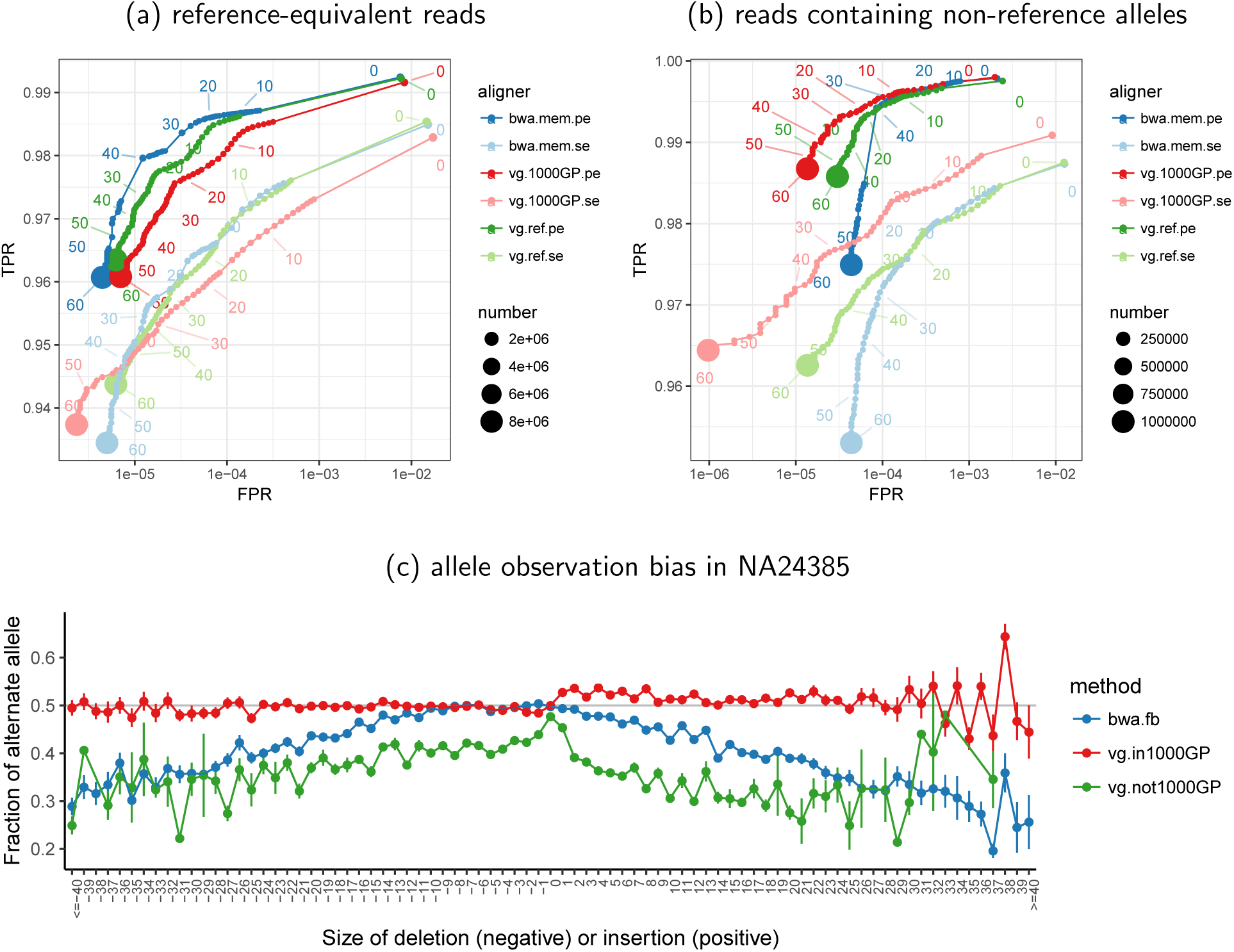
(a) the ROC curves for 10M read pairs simulated from NA24385 bwa mem, vg with a linear genome reference, and vg with the same pangenome reference. Performance is shown for both single end (se) and pair end (pe) mapping. (b) the alternate allele fraction at heterozygous variants called by GIAB in NA24385 as a function of deletion or insertion size. (c) as in (b) but as a function of distance to the nearest non-reference variant.

For the reads from reference-equivalent segments, both single end and paired end alignment with vg against the reference graph performed comparably to bwa mem (green and blue curves). Mapping these reads against the 1000GP variation graph (red curves) gave slightly worse performance over most of the range, which we attribute to a combination of the increase in options for alternative places to map reads provided by the variation graph, and the fact that we needed to prune some GCSA2 index k-mers in the most complex regions of the graph.

For reads that were simulated from segments containing non-reference alleles (approximately 10% of reads), vg mapping to the 1000GP graph gives better performance than either vg or bwa mem mapping to the linear reference (Figure 2b), due to the fact that many variants present in NA24385 are already represented in the 1000GP graph. This is particularly clear for single end mapping, since many paired end reads are rescued by the mate read mapping.

We also mapped a real human genome read set with approximately 50X coverage of Illumina 150bp paired end reads from the NA24385 sample to the 1000GP graph. vg produced mappings for 98.7% of the reads, 88.7% with reported mapping quality score ≥ 30 on the Phred scale, and 76.8% with perfect, full-length sequence identity to the reported path on the graph.

For comparison, we also used vg to map these reads to the linear reference. Similar proportions of reads mapped (98.7%) and mapped with reported quality ≥ 30 (88.8%), but considerably fewer with perfect identity (67.6%). Markedly different mappings were found for 1.0% of reads (0.9% mapping to widely separated positions on the two graphs, and 0.1% mapping to one graph but not the other). The reads mapping to widely separated positions were strongly enriched for repetitive DNA. For example, the linear reference mappings for 27.5% of these read pairs overlap various types of satellite DNA identified by RepeatMasker, compared to 3.0% of all read pairs.

To illustrate the consequences of mapping to a reference graph rather than a linear reference, we stratified the sites called as heterozygous insertions or deletions in NA24385 by the GIAB by deletion or insertion length (0 for single nucleotide variants) and by whether the site was present in 1000GP, and measured the fraction of reads mapped to the alternate allele for each category. The results show that mapping with vg to the population graph when the variant is present in 1000GP (95.4% of sites) gives nearly balanced coverage of alternate and reference alleles independent of variant size, whereas mapping to the linear reference either with vg or bwa mem leads to a progressively increasing bias with increasing deletion and (especially) insertion length (Figure 2c), so that for insertions around 30bp a majority of insertion containing reads are missing (there are over twice as many reference reads as alternate reads).

### 3.2 Mapping against non-ordered reference graphs

The graphs that we have used so far were constructed from variation data obtained from mapping to a linear reference, and so are directed acyclic graphs. We next demonstrate the ability of vg to work with arbitrary graphs that include duplications, inversions, and translocations, by showing its use with multiple yeast strains independently assembled *de novo* using long read data.^18^ These assemblies manifest large scale structural variation and novel sequence not detected in reference-based sequencing,^29^ including extensive rearrangement and reordering in subtelomeric regions^18^ as illustrated in Figure 1.

We compared four vg graphs: a linear reference graph from the standard S288c strain, a linear reference from the SK1 strain, a pangenome graph of all seven strains, and a “drop_SK1” variation graph in which all sequence private to the strain SK1 was removed from the pangenome graph. The multiple sample graphs were constructed with the Cactus progressive aligner,^6^ which non-partially ordered (directed-acyclic) graphs.

Similarly to the human experiments, we simulated 1 million 150bp paired reads from the SK1 reference, modelling sequencing errors, and mapped them to the four references. The resulting ROC curves are shown in Figure 3a. Not surprisingly, the best performance is obtained by mapping to a linear reference of the SK1 strain from which the data were simulated, with substantially higher sensitivity and specificity compared to mapping to the standard linear reference from the strain S288c with either vg or bwa mem. Mapping to the variation graphs gives intermediate performance, with several percent more sensitivity and lower false positive rates than to the standard reference. There is surprisingly little difference between mapping to graphs with and without the SK1 private variation, probably because much of what is novel in SK1 compared to the reference is also seen in other strains. Again we see lower sensitivity compared to mapping just to the SK1 sequence, likely because of suppression of GCSA2 index kmers in complex or duplicated regions.

**Figure 3:**
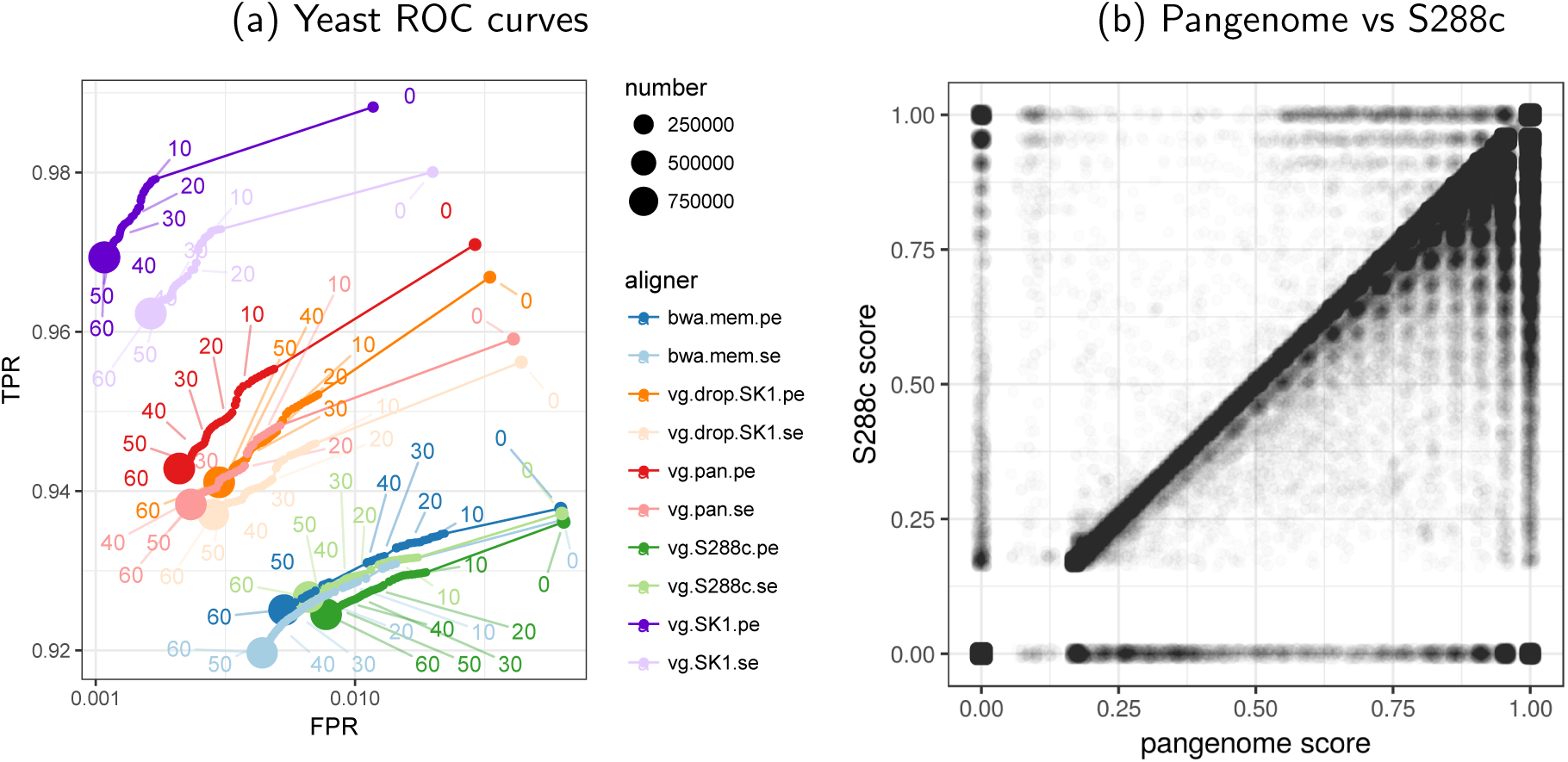
(a) ROC curves as in Figure 2a,b obtained by mapping simulated SK1 yeast strain reads against a variety of references described in the text; (b) a dotplot of alignment score relative to the maximum possible when mapping reads from 12 additional yeast strains against either the 7 strain pangenome variation graph or the S288c linear reference.

To demonstrate the difference between mapping to the pangenome and to the standard reference with real data, we downloaded from the National Centre for Yeast Cultures (NCYC, http://opendata.ifr.ac.uk/NCYC/) twelve 101bp Illumina paired end read sets for *S*. *cerevisiae* strains that were not included in the pangenome variation graph, and mapped them using
vg to the pangenome and S288c linear references. For this analysis we added the mitochondrial and 2 micron plasmid circular genomes to the pangenome. Slightly more reads map to the pangenome (93.7%) than to the linear reference (93.2%), but while 69.6% of reads map perfectly to the pangenome, only 52.8% map perfectly to the S288c linear reference. Of the reads that map, 74.6% map with equal scores to both the pangenome and linear references, 24.9% map better to the pangenome, and only 0.5% map better to the linear reference. Figure 3b shows a dotplot of the normalized alignment scores of reads to these two references, reflecting that much of the variation in these novel genomes could be found in even a small number of alternative reference sequences.

Finally, we constructed a vg graph from a *de novo* assembly of the cichlid fish *Astatotilapia calliptera* made with Falcon^30^ from Pacific Bioscience long read data, with 270,735 overlapping reads totalling 1.02Gbp after removing overlaps. We mapped to this an independent 81x depth Illumina data set, achieving 95.2% mapping rate.

## 4 Discussion

In conclusion, vg implements a suite of tools for genomic sequence data analysis using general variation graph references. Using the vg toolkit, we can construct or import a graph, modify it, visualize it, and use it as a reference. vg can accurately map new sequence reads to the reference using succinct indexes of the graph and its sequence space, and can describe variation between a new sample and an arbitrary reference embedded as a path in the graph. Elsewhere^17^ we discuss the use of vg to map read sets and call variants against a number of alternative human reference graphs built from multiple regions of the human genome with different properties.

For applications such as mapping to the 1000 Genomes Project variant call set, the recently published GraphTyper^12^ package also supports mapping to a graph. However, this functionality is limited compared to what is available in vg in two important ways. First, GraphTyper only locally realigns reads that already map, to the region they were already mapped to by a standard linear genome mapper such as bwa mem, rather than globally mapping to the graph as we do with our GCSA2 index. Second, it does not support arbitrary graph structures, which would be necessary to represent complex genome rearrangements (as in our yeast example) and assembly graphs.

There are many areas for potential future development and application of vg. These include further improvements in the mapping and variant calling algorithms, potentially using long range statistical haplotype structure information, stored in a graph extension of the PBWT haplotype compression and search data structure,^31^ as proposed by Novak.^32^ Beyond variant calling, the ability to map in an unbiased way to both reference and alternate alleles is potentially important when quantitating allele-specific expression^33^ or protein binding. We note that graphs can also naturally represent the relationships between transcribed, spliced and edited RNA sequences and the genome from which they are transcribed, so the vg software can potentially be used for splice-aware RNAseq mapping.^34^

We believe that genome variation graphs will underpin a new paradigm for genome sequence data analysis.^1^ They support the representation of structural variation using the same components (edges, nodes and paths) that are used to represent single base changes. For human, they allow more accurate and complete read mapping, as demonstrated in Figure 2. The benefits will only be greater for other organisms with higher levels of genetic variation, or for which uncertainties remain in the reference assembly. In order for the biological research community to exploit these advantages, it needs software for variation graphs that scales to the genomes of humans and other complex organisms. vg is a robust and openly available platform to fulfill this need.

## 5 Acknowledgements

EG, JS and RD were funded by the Wellcome Trust (grant 090851). ETD was funded by an NIH Cambridge Trust studentship, and WJ by a Wellcome Trust MGM studentship (109083/Z/15/Z). AN, GH, JE and BP were supported by the National Institutes of Health (5U41HG007234), the W. M. Keck Foundation (DT06172015) and the Simons Foundation (SFLIFE# 35190). MFL is employed by DNAnexus Inc. We thank members of the GA4GH Reference Variation Working Group for support, ideas and comments.

## 6 Author contributions

EG conceived and led the development of vg, JS developed the GCSA2 index, AMN, GH, JME and ETD contributed to the software, EG, WJ and ML contributed results and data analysis, BP and RD oversaw the project, and all contributed to the manuscript.

## S1 Supplement

### S1.1 GCSA2 index generation

We generate the GCSA2 index for a vg graph by transforming the graph into an effective De Bruijn graph with *k* = 64 or *k* = 128, with positional labels that indicate where in the source graph a particular kmer starts. GCSA2 can then generate a GCSA from this labeled De Bruijn graph. To build the De Bruijn graph efficiently, we initially generate kmers with *k* =16 labelled by their start, end, preceding and following bases in the vg graph. GCSA2 then undertakes two or three rounds of prefix doubling, wherein it uses the positional information in the input De Bruijn graph to increase the order of the graph.

In certain circumstances where there are repeated patterns of dense variants in the graph, the path complexity explodes exponentially, preventing practical enumeration of the De Bruijn graph. To avoid this, we prune edges from the vg graph which induce more than a certain number (default 4) of combinatorial bifurcations within a certain linear distance (default 16 bases), and remove any extremely short subgraphs that result from this destructive masking operation. This transformation preserves the coordinate system of the graph, allowing us to use it as the basis for seed generation against the unpruned graph.

### S1.2 SMEM and secondary MEM generation

vg can efficiently align reads against large graphs supported by xg and GCSA2 indexes through an alignment process based on maximal exact match sequences between the query sequence and the reference graph. First, a set of super-maximal exact matches (SMEMs) of a read are generated by traversing the suffix tree encoded in the GCSA2 index until the count of matching strings drops to 0, then backing off one step to find all longest exact matches. A secondary “reseed” pass through the traversal can then identify next-longest matches, which are used both to improve sensitivity and to evaluate mapping quality (see below). Then, chains of SMEMs that are consistent with the query sequence are found using a Markov model in which the optimal alignment is likely to form a Viterbi path. For each candidate chain, we then locally align the read against the graph. Scoring results from the local alignment are used to rank the candidate alignments. We then return the best alignment, or multiple candidates if multiple mappings are required.

### S1.3 SMEM chaining and read pairing

If the MEMs do not cover the full read length then we attempt to link them together into chains by building a Markov model in which the best possible chains form high-scoring paths. In this model the nodes correspond to the reference graph positions where MEMs in the read occur and the transitions between nodes correspond to a weight that is proportional to the approximate indel size implied by the approximate positions of the MEMs in the graph. If we are aligning a read pair, the weight between MEMs on different fragments is proportional to the probability of that distance under a learned model of the fragment distribution. One we establish this model, we take the Viterbi path through it as our first candidate alignment. By masking this path out and re-running the Viterbi algorithm on the model, we can extract a series of candidate alignments in descending order of goodness.

### S1.4 Matching SMEM chains to the graph

Given a candidate cluster of exact matches that we have extracted from the SMEM chaining model described in S1.3, we want to derive a complete description of the alignment of the read to the reference graph. Sensitive alignment of each candidate is essential for distinguishing the optimal alignment.

### S1.5 Banded alignment

For very long reads, where in the worst case the local dynamic programming can become prohibitively expensive, we break the reads into “bands” of a fixed width *w* (default 256 base pairs) with overlap between successive bands of *w*/2. We then align these bands independently, trim the overlaps from the alignments, and concatenate the results. This allows vg to map noisy reads of arbitrary length, and is used as a core component in the long read progressive assembler vg msga.

### S1.6 SIMD-accelerated local alignment

We link seeds using local alignment based on dynamic programming. To make this efficient we extended an implementation (Zhao et al., 2013) of Farrar’s SIMD-accelerated striped Smith Waterman (SSW) algorithm (Farrar, 2007), which we term “graph striped Smith-Waterman” GSSW. (Single input multiple data (SIMD) instructions allow vectorized mathematical operations in a single machine instruction, and can be used to greatly speed up algorithms which can be implemented in terms of operations on vectors.) GSSW generalizes all aspects of SSW to operate over sequence directed acyclic graphs, including affine gap penalties, and retains its matrices for traceback.

To interface with GSSW we transform a local region of our graph so that it is acyclic and only represents a single strand of the DNA. The two operations we use in this transformation are *unfold,* which expands the graph to include its reverse complement where accessible via an inversion, and *dagify,* which unrolls strongly connected components of the graph far enough that we are guaranteed to be able to find any sequence of given length *k* in the source graph in the unrolled one. This allows us to align any sequence of up to length *k* against a completely general section of a variation graph. Through these steps we retain a mapping from old node ids to new ones, which we will use to project alignments to the transformed graph back into our base coordinate space.

### S1.7 Unfolding

Every node has an implicit default orientation (see Graph Representation below) so that it is possible to determine edges that cause an inversion, i.e. those which connect between a forward and a reverse complement node orientation. In VG::unfold we use a breadth first search starting at every inverting edge in the graph to explore the reverse complemented portions of the graph that we can reach within length k from the inverting edge. We then copy this subgraph, take its reverse complement, and replace the inverting edges connecting it to the forward strand of the graph with non-inverting ones. If k is greater than any length in our graph, then we are guaranteed to duplicate the entire reverse complement of the graph on the forward strand, effectively doubling the size of the graph if we have any inversions in it, as shown in Figure S1.

### S1.8 Dagification

Variation graphs may have cycles. These are useful as compact representations of copy number variable regions, and arise naturally in the process of genome assembly. However, our partial order alignment implementation does not handle these structures, and so when they occur we convert them into an approximately equivalent acyclic graph in order to align with GSSW. To do so, we unroll cyclic structures by copying their internal nodes an appropriate number of times to allow a given query length to align through the unrolled version of the component.

We first detect all strongly connected components by using a recursion-free implementation of Tarjan’s strongly connected components algorithm (Tarjan, 1972). Then, we copy each strongly connected component and its internal edges into a new graph. We greedily break edges in this graph that introduce cycles. Next we k-DAGify the component progressively copying the base component and, for each edge between nodes in the component, connecting from the source node in the previous copy to the target node in the current copy.

We use dynamic programming to track the minimum distance back through the graph to a root node outside the component at each step. When this reaches our target *k*, we stop unrolling, and add the expanded component back into the graph by reconnecting it with its original neighborhood. For each copy of a node in the DAG-ified component we copy all its inbound and outbound edges where the other end of the edge lies outside the strongly connected component. The resulting graph is acyclic and supports queries up to length k on the original graph using the translation that we maintain between the new graph and the source one. Figure S2 provides a visual description of the process.

### S1.9 Base quality adjusted alignment scores and mapping qualities

Base qualities are typically reported on the Phred scale so that the probability of error for a given quality *Q* is ∈ = 10^-*Q*/10^. Assuming no bias in which bases are mistaken for each other, this defines a posterior distribution over bases *b* for a base call *x*.

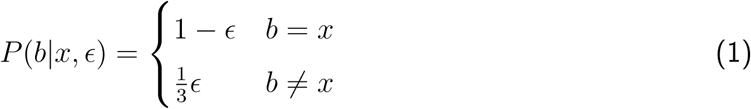

We use this distribution to derive an adjusted score function. Normally, the match score for two bases is defined as the logarithm of the likelihood ratio between seeing two bases *x* and *y* aligned and seeing them occur at random according to their background frequencies.

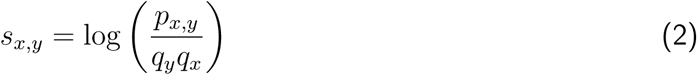

Next we marginalize over bases from the posterior distribution to obtain a quality adjusted match score.

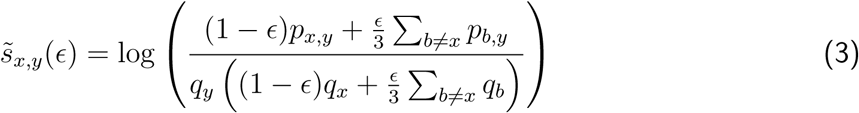

vg works backwards from integer scoring functions to the probabilistic alignment parameters in this equation. After doing so, the match scores are given by

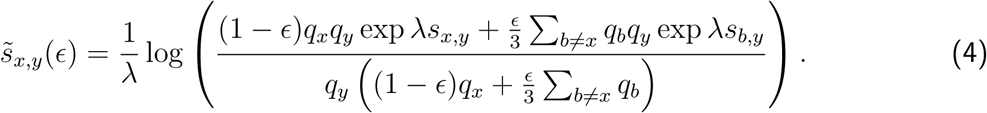

Here, λ is a scale factor that can be computed from the scoring parameters, and the background frequencies *q_x_* are estimated by their frequency in the reference graph. Since base quality scores are already discretized, the adjusted scores can be precomputed and cached for all reasonable values of ∈.

### S1.1. Mapping qualities

The algorithm for mapping qualities in vg is also motivated by a probabilistic interpretation of alignment scores. The score of an alignment A of two sequences X and Y is the sum of scores given in (2). This makes it a logarithm of a joint likelihood ratio across bases, where the bases are assumed independent (a more complete justification including gap penalties involves a hidden Markov model, but it can be shown to approximate this formula). We denote this score *S*(*A|X, Y*). Thus, assuming a uniform prior over alignments, we can use Bayes’ Rule to motivate a formula for the Phred scaled quality of the optimal alignment, 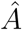.

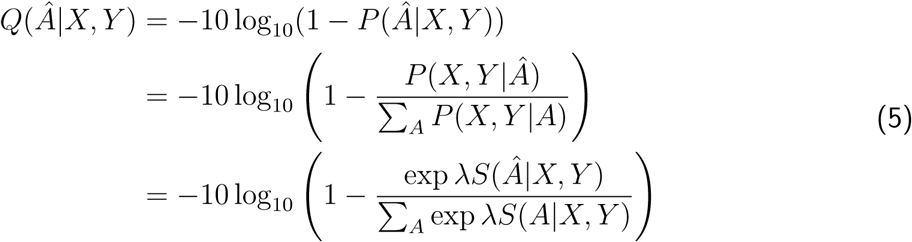

Using the close approximation of the LogSumExp function by element-wise maximum, there is a fast approximation to this formula that does not involve transcendental functions.

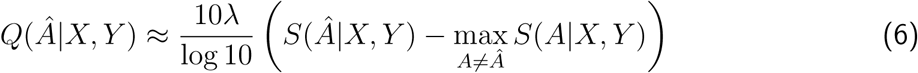

In practice, we do not compare the optimal alignment to all possible alignments, but to the optimal alignments from other seeds. Thus, the mapping quality indicates the confidence that we have aligned the read to approximately the correct part of the graph rather then that the fine-grained alignment in that part of the graph is correct. It is also worth noting that since this formula is based on alignment scores, it can incorporate base quality information through the base quality adjusted alignment scores.

### S1.11 Succinct graph representation (xg)

When using a variation graph as a reference system, we are unlikely to need to modify it. As such we can compress it into a system that provides efficient access to important aspects of the graph. Specifically, we care about the node and edge structure of the graph and queries that allow us to extract and seek to positions in embedded paths. We would like to be able to query a part of the graph corresponding to a particular region of a chromosome in a reference path embedded in the graph. Similarly, if we find an exact match on the graph using GCSA2, we would like to load that region of the graph into memory for efficient local alignment.

We implement a succinct representation of variation graphs in the xg library, using data structures from SDSL-lite. Node labels and node ids are stored in a collection of succinct vectors, augmented by rank/select dictionaries that allow the lookup of node sequences and node ids. An internal node rank is given for each node, and we map from and to this internal coordinate system using a compressed integer vector of the same order as the node id range of the graph we have indexed. To allow efficient exploration of the graph, we store each node’s edge context in a structured manner in an integer vector, into which we can jump via a rank/select dictionary keyed by node rank in the graph. Efficient traversal of the graph’s topology via this structure is enabled by storing edges a relative offsets to “to” or “from” node, which obviates the need for secondary lookups and reduces the cost of step-wise traversal to member access on the containing vector and the cost of parsing each node context record that we encounter. Paths provided to XG are used to induce multifarious coordinate systems over the graph. We store them using a collection of integer vectors and rank/select dictionaries that allow for efficient queries of the paths at or near a given graph position, as well as queries that give us the graph context near a given path position.

---

**Figure S1:**
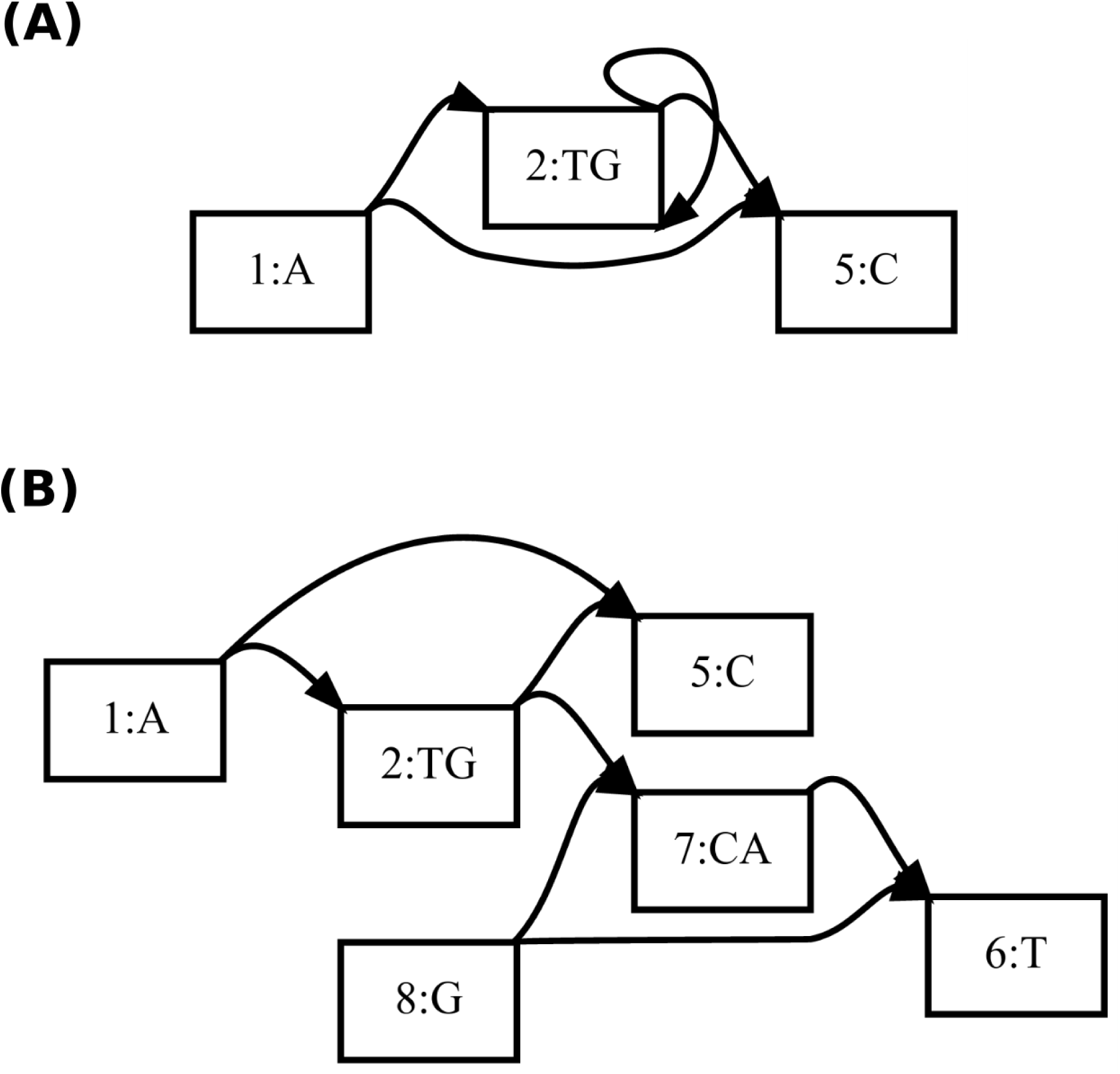
Illustration of the unfolding process. The starting graph (A) has an inverting edge leading from the forward to reverse strand of node 2. In (B) we unroll the graph with k greater than the length of the graph, which materializes the implied reverse strand as sequence on the forward strand of new nodes.

**Figure S2:**
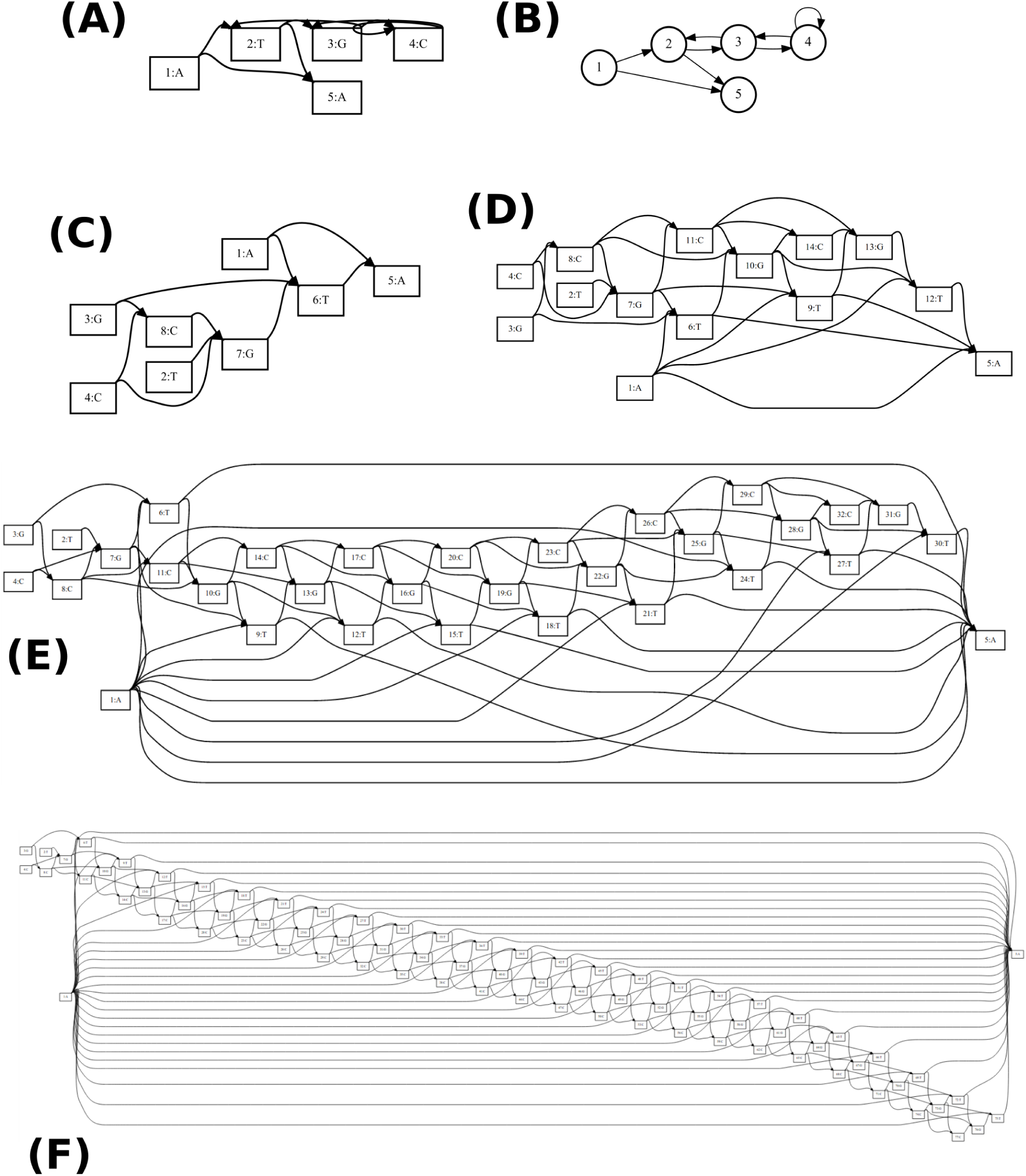
Illustration of the unrolling process. The starting graph (A) and a representation without sequences or sides to clarify the underlying structure (B). In (C) we have unrolled one step (*k* = 2). In (D), *k* = 4, (E) *k =* 10, and (F) *k =* 25.

## References

1 B. Paten, A. M. Novak, J. M. Eizenga, and E. Garrison. Genome graphs and the evolution of genome inference. Genome Res., 27(5):665–676, 2017.

2 Pavel A Pevzner, Haixu Tang, and Michael S Waterman. An eulerian path approach to dna fragment assembly. Proceedings of the National Academy of Sciences, 98(17):9748–9753, 2001.

3 E. W. Myers. The fragment assembly string graph. Bioinformatics, 21 Suppl 2:79–85, Sep 2005.

4 Jouni Sirén. Indexing variation graphs. In 2017 Proceedings of the ninteenth workshop on algorithm engineering and experiments (ALENEX), pages 13–27. SIAM, 2017.

5 A. L. Delcher, S. Kasif, R. D. Fleischmann, J. Peterson, O. White, and S. L. Salzberg. Alignment of whole genomes. Nucleic Acids Res., 27(11):2369–2376, Jun 1999.

6 Benedict Paten, Dent Earl, Ngan Nguyen, Mark Diekhans, Daniel Zerbino, and David Haussler. Cactus: Algorithms for genome multiple sequence alignment. Genome Research, 21(9):1512–1528, September 2011.

7 Ruiqiang Li, Yingrui Li, Hancheng Zheng, Ruibang Luo, Hongmei Zhu, Qibin Li, Wubin Qian, Yuanyuan Ren, Geng Tian, Jinxiang Li, et al. Building the sequence map of the human pangenome. Nature biotechnology, 28(1):57–63, 2010.

8 Shuai Yuan and Zhaohui Qin. Read-mapping using personalized diploid reference genome for rna sequencing data reduced bias for detecting allele-specific expression. 2012 IEEE International Conference on Bioinformatics and Biomedicine Workshops, Oct 2012.

9 C. Lee, C. Grasso, and M. F. Sharlow. Multiple sequence alignment using partial order graphs. Bioinformatics, 18(3):452–464, Mar 2002.

10 Petr Danecek, Adam Auton, Goncalo Abecasis, Cornelis A Albers, Eric Banks, Mark A DePristo, Robert E Handsaker, Gerton Lunter, Gabor T Marth, Stephen T Sherry, et al. The variant call format and VCFtools. Bioinformatics, 27(15):2156–2158, 2011.

11 A. Dilthey, C. Cox, Z. Iqbal, M. R. Nelson, and G. McVean. Improved genome inference in the MHC using a population reference graph. Nat. Genet., 47(6):682–688, Jun 2015.

12 Hannes P Eggertsson, Hakon Jonsson, Snaedis Kristmundsdottir, Eirikur Hjartarson, Birte Kehr, Gisli Masson, Florian Zink, Kristjan E Hjorleifsson, Aslaug Jonasdottir, Adalbjorg Jonasdottir, et al. Graphtyper enables population-scale genotyping using pangenome graphs. Technical report, Nature Research, 2017.

13 Goran Rakocevic, Vladimir Semenyuk, James Spencer, John Browning, Ivan Johnson, Vladan Arsenijevic, Jelena Nadj, Kaushik Ghose, Maria C Suciu, Sun-Gou Ji, et al. Fast and accurate genomic analyses using genome graphs. bioRxiv, page 194530, 2017.

14 Erik Garrison and Gabor Marth. Haplotype-based variant detection from short-read sequencing. arXiv preprint arXiv:1207.3907, 2012.

15 M. A. DePristo, E. Banks, R. Poplin, K. V. Garimella, J. R. Maguire, C. Hartl, A. A. Philippakis, G. del Angel, M. A. Rivas, M. Hanna, A. McKenna, T. J. Fennell, A. M. Kernytsky, A. Y. Sivachenko, K. Cibulskis, S. B. Gabriel, D. Altshuler, and M. J. Daly. A framework for variation discovery and genotyping using next-generation DNA sequencing data. Nat. Genet., 43(5):491–498, May 2011.

16 1000 Genomes Project Consortium et al. A global reference for human genetic variation. Nature, 526(7571):68–74, 2015.

17 Adam M Novak, Glenn Hickey, Erik Garrison, Sean Blum, Abram Connelly, Alexander Dilthey, Jordan Eizenga, MA Saleh Elmohamed, Sally Guthrie, André Kahles, et al. Genome graphs. bioRxiv, page 101378, 2017.

18 Jia-Xing Yue, Jing Li, Louise Aigrain, Johan Hallin, Karl Persson, Karen Oliver, Anders Bergström, Paul Coupland, Jonas Warringer, Marco Cosentino Lagomarsino, et al. Contrasting evolutionary genome dynamics between domesticated and wild yeasts. Nature genetics, 49(6):913–924, 2017.

19 H. Li, B. Handsaker, A. Wysoker, T. Fennell, J. Ruan, N. Homer, G. Marth, G. Abecasis, and R. Durbin. The Sequence Alignment/Map format and SAMtools. Bioinformatics, 25(16):2078–2079, Aug 2009.

20 Simon Gog, Timo Beller, Alistair Moffat, and Matthias Petri. From theory to practice: Plug and play with succinct data structures. In International Symposium on Experimental Algorithms, pages 326–337. Springer, 2014.

21 Ora Lassila and Ralph R Swick. Resource description framework (rdf) model and syntax specification. 1999.

22 E. W. Myers and W. Miller. Approximate matching of regular expressions. Bull. Math. Biol., 51(1):5–37, 1989.

23 M. Farrar. Striped Smith-Waterman speeds database searches six times over other SIMD implementations. Bioinformatics, 23(2):156–161, Jan 2007.

24 Richard Durbin, Sean R Eddy, Anders Krogh, and Graeme Mitchison. Biological sequence analysis: probabilistic models of proteins and nucleic acids. Cambridge University Press, 1998.

25 Michiaki Hamada, Edward Wijaya, Martin C Frith, and Kiyoshi Asai. Probabilistic alignments with quality scores: an application to short-read mapping toward accurate snp/indel detection. Bioinformatics, 27(22):3085–3092, 2011.

26 H. Li. A statistical framework for SNP calling, mutation discovery, association mapping and population genetical parameter estimation from sequencing data. Bioinformatics, 27(21):2987–2993, Nov 2011.

27 Heng Li. Aligning sequence reads, clone sequences and assembly contigs with bwa-mem. arXiv preprint arXiv:1303.3997, 2013.

28 Justin M Zook, David Catoe, Jennifer McDaniel, Lindsay Vang, Noah Spies, Arend Sidow, Ziming Weng, Yuling Liu, Christopher E Mason, Noah Alexander, et al. Extensive sequencing of seven human genomes to characterize benchmark reference materials. Scientific data, 3, 2016.

29 G. Liti, D. M. Carter, A. M. Moses, J. Warringer, L. Parts, S. A. James, R. P. Davey, I. N. Roberts, A. Burt, V. Koufopanou, I. J. Tsai, C. M. Bergman, D. Bensasson, M. J. O’Kelly, A. van Oudenaarden, D. B. Barton, E. Bailes, A. N. Nguyen, M. Jones, M. A. Quail, I. Goodhead, S. Sims, F. Smith, A. Blomberg, R. Durbin, and E. J. Louis. Population genomics of domestic and wild yeasts. Nature, 458(7236):337–341, Mar 2009.

30 Chen-Shan Chin, Paul Peluso, Fritz J Sedlazeck, Maria Nattestad, Gregory T Concepcion, Alicia Clum, Christopher Dunn, Ronan O’Malley, Rosa Figueroa-Balderas, Abraham Morales-Cruz, et al. Phased diploid genome assembly with single-molecule real-time sequencing. Nature methods, 13(12):1050–1054, 2016.

31 Richard Durbin. Efficient haplotype matching and storage using the positional Burrows-Wheeler transform (PBWT). Bioinformatics, 30(9):1266–1272, 2014.

32 AM Novak, E Garrison, and B Paten. A graph extension of the positional Burrows-Wheeler transform and its applications. In M Firth and CN Pedersen, editors, Algorithms in bioinformatics, pages 246–256. Springer, Heidelberg, Germany, 2016.

33 B. Ge, D. K. Pokholok, T. Kwan, E. Grundberg, L. Morcos, D. J. Verlaan, J. Le, V. Koka, K. C. Lam, V. Gagne, J. Dias, R. Hoberman, A. Montpetit, M. M. Joly, E. J. Harvey, D. Sinnett, P. Beaulieu, R. Hamon, A. Graziani, K. Dewar, E. Harmsen, J. Majewski, H. H. Goring, A. K. Naumova, M. Blanchette, K. L. Gunderson, and T. Pastinen. Global patterns of cis variation in human cells revealed by high-density allelic expression analysis. Nat. Genet., 41(11):1216–1222, Nov 2009.

34 S. Beretta, P. Bonizzoni, L. Denti, M. Previtali, and Rizzi R. Mapping RNA-seq Data to a Transcript Graph via Approximate Pattern Matching to a Hypertext. In D. Figueiredo, C. Martín-Vide, D. Pratas, and M. Vega-Rodriguez, editors, Algorithms for Computational Biology. AlCoB 2017, pages 49–61. Lecture Notes in Computer Science, vol 10252. Springer, Cham, 2017.

## References

Farrar, M. Striped Smith-Waterman database searches six times over other SIMD implementations. Bioinformatics, 23(2):156–161, 2007.

Grossi, R., Gupta, A. and Scott Vitter, J. High-order entropy-compressed text indices. In Proceedings of the Fourteenth Annual ACM-SIAM Symposium on Discrete Algorithms, pages 841–850, Society for Industrial and Applied Mathematics, 2003.

Okanohara, D. and Sadakane, K. Practical entropy-compressed rank/select dictionary. In Proceedings of the Meeting on Algorithm Engineering & Experiments, pages 60–70. Society for Industrial and Applied Mathematics, 2007.

Tarjan, R. Depth-first search and linear graph algorithms. SIAM Journal on Computing. 1(2):146–160, 1972.

Zhao, M., Lee, W-P., Garrison, E. and Marth, G. SSW library: An SIMD smith-waterman C/C++ library for use in genomic applications. PloS One, 8:e82138, 2013.

